# Cell Chromatography: Biocompatible chromatographic separation and interrogation of microbial cells

**DOI:** 10.1101/2022.03.07.482710

**Authors:** M. Hazu, A. Ahmed, E. Curry, D.P. Hornby, D.T. Gjerde

## Abstract

The isolation of pure, single colonies lies at the heart of experimental microbiology. However, a microbial colony typically contains around one million cells at all stages of the life cycle. Here we describe a novel, cell chromatography method that facilitates the capture, purification and interrogation of microbial cells from both single and mixed cultures. The method described relies on, but is not limited to, differences in surface charge to separate bacterial strains. The method is fully biocompatible, leading to no significant loss of cell viability,. The chromatographic capture of cells, combined with selective elution methods facilitates a greater level of experimental control over the sample inputs required for downstream high throughput and high sensitivity, analytical methods. The application of the method for interrogating the antibiotic resistance of bacterial strains and for the separation of bacteria from environmental samples is illustrated.

## Introduction

The isolation of single colonies using selective growth media has been the mainstay of microbiology since the development of the Petri dish over 100 years ago. However, each colony of any pure bacterial strain is intrinsically heterogeneous with respect to its physiological growth status. In most cases, single colonies are initially obtained by subculturing in the laboratory from a Petri dish or broth culture, or from “the field”, via a clinical or environmental sample. Following some form of streaking or spreading protocol, combined with an empirical dilution of the sample, single colonies are typically obtained overnight (depending on the growth kinetics of the strain under investigation and the suitability of the growth medium). A single colony will typically contain around 1 million cells, all of which will be at a different stage in the growth cycle: at extremes, some will be newly replicated while others will be dead. However, as molecular analysis techniques become increasingly powerful, higher in throughput and capable of single cell resolution, the traditional methods of microbiology for both the analytical and preparative purification of cells have become limiting.

Liquid chromatography has been one of the most successful and enduring methods for the purification of biological molecules since its widespread uptake by biochemists over the last 60 years (see for example Duong-Ly and Gabelli (2014). Many commercial suppliers of chromatography media and instruments have focused on the separation of proteins, peptides and nucleic acids, where a wide range of protocols and chromatographic media have been developed which combine high resolution with retention of biochemical function. In parallel, chromatography resins and instruments have been developed to support downstream chemical analysis (mainly peptide sequence determination) where functional integrity is not required. Mass spectrometry of proteins and peptides is probably the best example of a technology that is dependent on the chemical, but not functional integrity of input samples. In contrast, biochemical assays and structural biology techniques including X-ray crystallography, NMR spectroscopy and cryo-electron microscopy, all demand both high levels of purity and functional integrity of the sample.

As microscopic techniques for interrogating cells and tissues approach the resolution of macromolecular structure determination methods (reviewed in Thorn, 2016), the demand for reproducible cell purification methods is becoming increasingly important. The first successful application of liquid chromatography for the separation of microbial cells has been described by Arvidsson et al (2002), who used super-macroporous cryogels for the separation of bacteria. They have shown that ion exchange and immobilised metal ion chromatography can be used to separate microbial cell mixtures with moderate retention of cell viability. Using conventional liquid chromatography protocols, bacterial cells are applied to a cryogel column and are captured at low ionic strength. The captured cells are subsequently eluted following methods that are commonly applied to protein separation.

Here we describe a novel method for the purification of cells using ion exchange chromatography, although there is no *a priori* limit to the choice of stationary phase medium. However, unlike conventional chromatography, cells and buffers enter and exit the column in a back-and-forth flow (Gjerde, 2018, 2019). We refer to this method simply as “cell chromatography”. This method has been applied to the separation of a mixture of gram positive (*Staphylococcus aureus*) and gram negative (*Escherichia coli*) bacteria. In addition, we demonstrate the resolving power of the method for the separation of individual strains found in local environmental water samples. Just as proteins in complex cell extracts can be separated by ion exchange chromatography, the differences in net surface charge are sufficient to separate heterogeneous populations of cells, with no significant loss of viability.

## Materials and Methods

### Growth and isolation of bacterial strains

*Staphylococcus aureus* SH100 (a kind gift from Professor Simon Foster, The University of Sheffield) and *Escherichia coli* BL21(DE3) with or without prior transformation with the expression vector pET22b (Novagen), were cultured on Luria Broth (with or without agar) and supplemented with 200µg/ml ampicillin (Sigma) as appropriate. Environmental samples of pond water (10ml) were collected in sterile tubes from a local boating lake adjoining the University of Sheffield, and were used on the same day, with or without dilution into phosphate buffered saline (PBS) at pH7.4. All standard methods were carried out as described in Sambrook et al (1989).

### Chromatography

Pipette tip-based columns provided by Gjerde Technologies contained a strong base anion exchanger bound to a water swollen agarose substrate. The columns were packed without compression to retain flow paths that are non-restrictive to bacterial cells. Pipette tip columns containing 100µl of resin Cytiva Q Sepharose® Fast Flow anion exchanger 90 micron average particle size in a 1 mL pipette tip with 0.635cm diameter frits were placed firmly onto an automatic pipette ensuring a tight seal to allow for efficient and reproducible uptake of liquids. Each column was first equilibrated by withdrawing 500µl of PBS solution, pH 7.4, at a rate of 0.75ml/min from a 96 well plate or Eppendorf tube and then, after 10s, 400µl of the PBS solution was ejected. Care was taken to ensure the column did not dry out: the total volume of liquid in the column was maintained above 100µl. The total volume of the resin was ∼100µl. The tips were mounted on a 6-channel, semi-automatic pipette, and flow rates and directions were driven by automated pipette software. For the work performed here Rainin E4 pipettes equipped with Purespeed® software (Biotage, San Jose) controlled the back-and-forth flow.

Typical bacterial suspensions were generated containing between 10^4^-10^5^ cells/ml of Luria Broth. These cell densities could be reproducibly and conveniently counted on agar plates following elution. In a typical protocol, 500µl of a bacterial suspension was withdrawn form a liquid culture at a rate of 0.75ml/min bringing the total column volume to 600µl. After 10s, 500µl of the column contents were ejected at the same rate, releasing the majority of unbound bacteria

In order to remove any bacteria that had bound non-specifically to the column matrix or other areas of the column, such as the exterior of the tip, the column was washed with 25 consecutive column volumes LB broth. The bound fraction of cells was then eluted with 500µl of LB broth into an Eppendorf tube and the total volume adjusted to 1ml with LB broth (for immediate use) or LB supplemented with glycerol (final concentration 25% v/v) if samples were to be frozen. To prevent further growth during chromatography, samples were washed with sterile PBS, without any significant loss of cell viability.

Elution of bound cells was achieved as follows, after 25 consecutive LB (or PBS) washes, matrix bound cells were eluted by back-and-forth addition of either LB or PBS containing increasing concentrations of NaCl from 50mM to 1M. At each NaCl concentration, 500µl of LB broth (or PBS), containing the required concentration of NaCl, from a total volume of 1ml in an Eppendorf was applied at a rate of 0.75ml/min and then immediately ejected.

Fractions were either analysed immediately or frozen in order to minimise any further growth of the cells before plating. The whole process takes approximately two hours to complete, depending on sample volumes and the number of wash and elute cycles required.

Eluted bacterial fractions were then plated on LB agar (with or without ampicillin) and incubated for 16 hours at 37°C. Bacterial cell numbers (colony forming units, CFUs) were determined by serial dilution from broth cultures followed by agar plating. All experiments were carried out in triplicate unless otherwise indicated. The arrangement of the pipette tip chromatography columns is shown in Figure 1.

**Figure 1.**
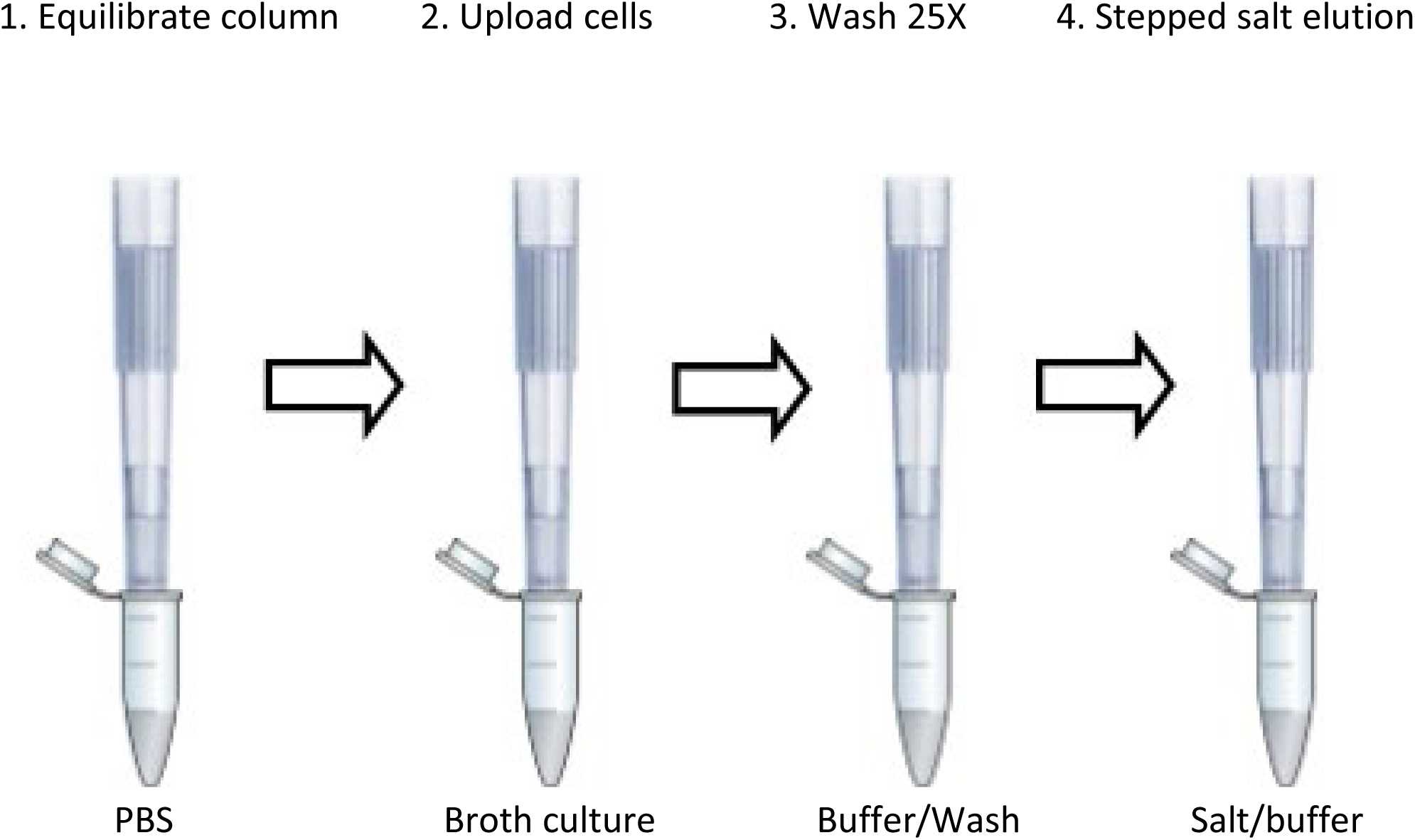
General illustration of the cell chromatography columns in a tip format, suitable for use with most commercial semi-automatic or fully automated, air-displacement pipetted. In step 1, the column is equilibrated by back and forth flow of the chosen buffer. In step 2, cells are “uploaded” from a broth culture or field sample, followed by extensive back and forth washing with loading buffer. Finally, in step 4, cells are eluted by step-wise additions of (in this case) PBS supplemented with NaCl.

## Results

### Quantifying capture and release of bacteria via cell chromatography

Throughout all of the experiments reported here a single protocol was used in which either broth cultures or liquid, field samples were applied using regulated back-and-forth flow through an ion exchange resin packed within conventional 1ml, disposable pipette tips.

The retention of viability of cells following cell chromatography is exemplified using the common laboratory strain, *E*.*coli* BL21 (DE3) in Table 1. There is a small reduction in colony forming units (CFUs) as a result of the separation, but this is most likely a reflection of small differences in external adherence of cells to the surface of the tips during manipulation.

**Table 1.**
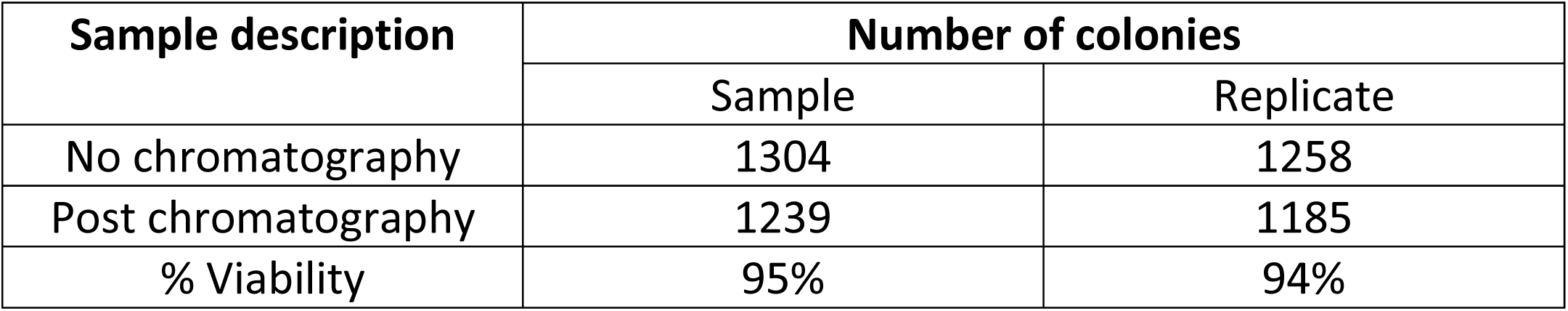
Estimation of the impact of cell chromatography on cell viability. Equivalent serial dilutions were made of cells before and after uploading and elution of *E*.*coli* BL21 (DE3) from two independent experiments. This protocol was incorporated as a control for each of the chromatography experiments described and the viability was never less than 90%.

Similar results were obtained for a number of other laboratory strains. The results in Table 2 show that approximately 5-10% of cells from a given broth culture are captured by the columns in the format supplied. In most experiments, the capacity of the columns is compatible with the requirements of downstream analysis, in this case using agar plates to determine the outcome of a particular experiment. The volume of chromatography medium can be increased (or decreased) to fit a particular application: for example, there was no significant difference in performance using larger bed volumes of up to several hundred µl (unpublished data).

**Table 2a.** Estimation of the fraction of uploaded *E. coli* cells bound to individual columns. 3 × 10^5 cells were uploaded in each case and the proportion of cells bound determined by serial dilution and standard cfu scoring on agar plates

| [NaCl] (mM) | Colonies |  |  |  |
| --- | --- | --- | --- | --- |
|  | Fraction 1 | Replicate 1 | Replicate 2 | Replicate 3 |
| 25<sup>th</sup> Wash Fraction (0mM) | 984 | 804 | 863 | 973 |
| 50 | 3704 | 1984 | 2345 | 2764 |
| 100 | 4128 | 3682 | 4032 | 4256 |
| 150 | 5728 | 4893 | 5409 | 5610 |
| 200 | 5936 | 4690 | 5537 | 6109 |
| 300 | 4902 | 5783 | 4723 | 5267 |
| 400 | 1928 | 1423 | 1746 | 2105 |
| 500 | 1112 | 812 | 1092 | 1273 |
| 750 | 416 | 444 | 321 | 492 |
| 1000 | 146 | 128 | 113 | 182 |
| TOTAL | 28000 | 23839 | 25318 | 28058 |
| % Cells bound | 9.33% | 7.95% | 8.44% | 9.35% |

**Table 2b.** Estimation of the fraction of uploaded *E. coli* cells bound to individual columns. 5 × 10^5 cells were uploaded in each case and the proportion of cells bound determined by serial dilution and standard cfu scoring on agar plates

| [NaCl] (mM) | Colonies |  |  |  |
| --- | --- | --- | --- | --- |
|  | Fraction 1 | Replicate 1 | Replicate 2 | Replicate 3 |
| 25<sup>th</sup> Wash Fraction (0mM) | 213 | 173 | 167 | 193 |
| 50 | 693 | 478 | 435 | 528 |
| 100 | 1363 | 1223 | 1290 | 1309 |
| 150 | 1765 | 1545 | 1368 | 1573 |
| 200 | 1492 | 1145 | 1092 | 1390 |
| 300 | 579 | 431 | 675 | 545 |
| 400 | 331 | 359 | 202 | 298 |
| 500 | 132 | 123 | 98 | 105 |
| 750 | 108 | 93 | 65 | 43 |
| 1000 | 27 | 83 | 40 | 7 |
| TOTAL | 6190 | 5478 | 5265 | 5798 |
| % Cells bound | 12.38% | 10.96% | 10.53% | 11.59% |

**Table 2c.** Estimation of the fraction of uploaded *S*.*aureus* cells bound to individual columns. 3 × 10^5 cells were uploaded in each case and the proportion of cells bound determined by serial dilution and standard cfu scoring on agar plates

| [NaCl] (mM) | Colonies |  |  |  |
| --- | --- | --- | --- | --- |
|  | Fraction 1 | Replicate 1 | Replicate 2 | Replicate 3 |
| 25<sup>th</sup> Wash<br>Fraction (0mM) | 444 | 509 | 549 | 465 |
| 50 | 2965 | 2305 | 2789 | 2340 |
| 100 | 3520 | 3109 | 3356 | 3271 |
| 150 | 4509 | 3756 | 4137 | 3987 |
| 200 | 3229 | 3912 | 3547 | 3730 |
| 300 | 1732 | 1907 | 1809 | 1869 |
| 400 | 902 | 933 | 810 | 758 |
| 500 | 630 | 739 | 548 | 586 |
| 750 | 214 | 312 | 253 | 201 |
| 1000 | 89 | 119 | 133 | 79 |
| TOTAL | 17790 | 17092 | 17382 | 16821 |
| % Cells bound | 5.93% | 5.70% | 5.79% | 5.61% |

**Table 2d.** Estimation of the fraction of uploaded *S*.*aureus* cells bound to individual columns. 5 × 10^5 cells were uploaded in each case and the proportion of cells bound determined by serial dilution and standard cfu scoring on agar plates.

| [NaCl] (mM) | Colonies |  |  |  |
| --- | --- | --- | --- | --- |
|  | Fraction 1 | Replicate 1 | Replicate 2 | Replicate 3 |
| 25<sup>th</sup> Wash Fraction (0mM) | 145 | 136 | 146 | 123 |
| 50 | 489 | 462 | 507 | 424 |
| 100 | 798 | 875 | 806 | 693 |
| 150 | 1232 | 1156 | 1178 | 1098 |
| 200 | 920 | 815 | 904 | 722 |
| 300 | 789 | 642 | 736 | 634 |
| 400 | 334 | 309 | 312 | 298 |
| 500 | 145 | 136 | 176 | 117 |
| 750 | 98 | 78 | 102 | 86 |
| 1000 | 46 | 67 | 56 | 60 |
| TOTAL | 4851 | 4540 | 4777 | 4132 |
| % Cells bound | 9.70% | 9.08% | 9.56% | 8.23% |

### Separation of bacterial cells by cell chromatography

The primary aim of this work was to determine whether the cell surface characteristics of living cells could be exploited to separate bacterial cells in a manner analogous to ion exchange chromatography of proteins. Separate broth cultures of *E. coli BL21* (carrying pET22b, an ampicillin resistant plasmid) and *S. aureus* (SH1000) were applied to ion exchange tips as described in the materials and methods section. The results of a step-wise application of NaCl as eluting agent are shown diagrammatically in Figure 2a. At pH7.4 *E*.*coli* and *S*.*aureus* strains can be adequately resolved, with some degree of overlap. Pure cultures can readily be obtained by taking early eluting fractions from the *E*.*coli* profile, or late eluting fractions from the *S*.*aureus* cells.

**Fig. 2a.**
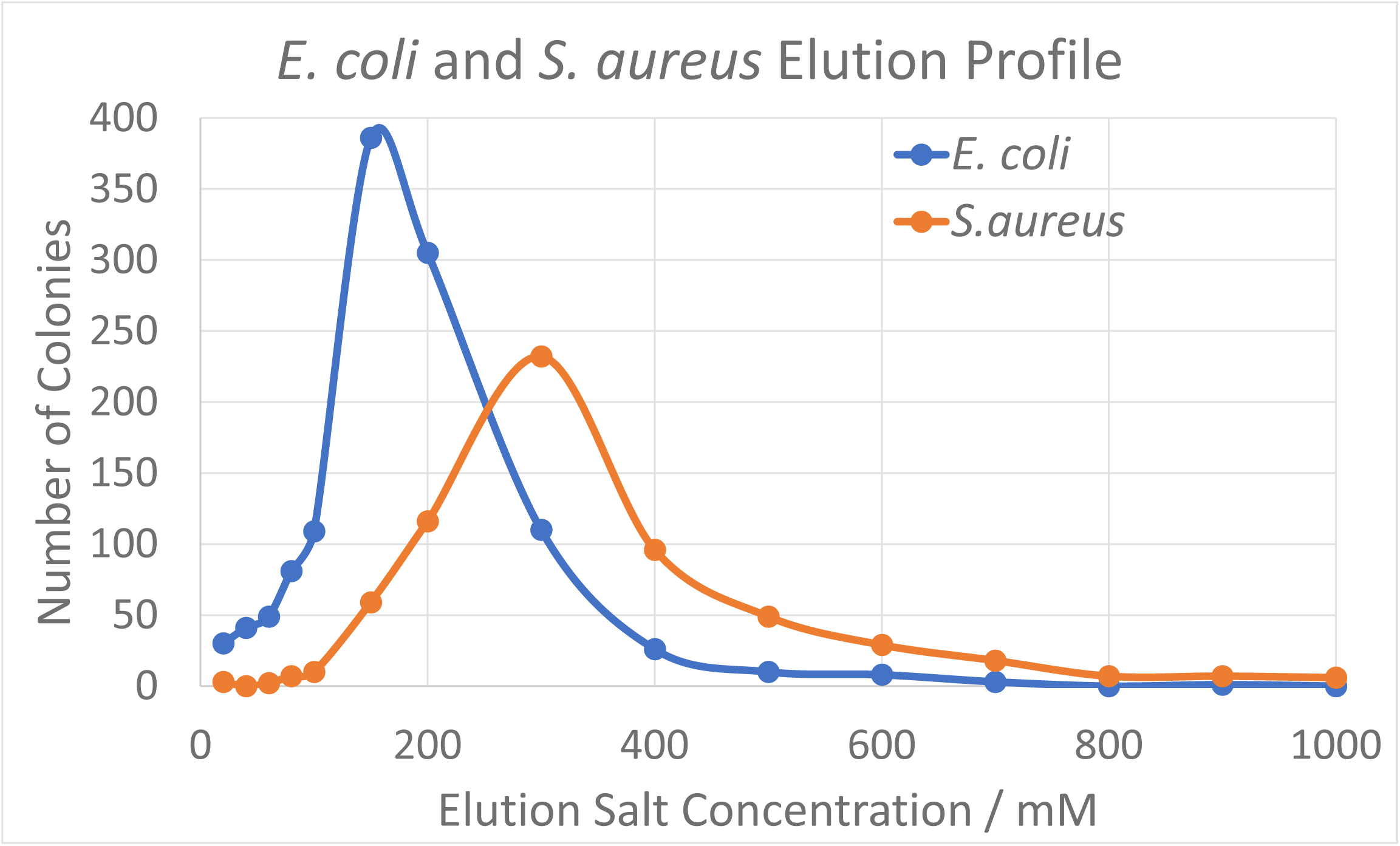
Separation of *E*.*coli* and *S. aureus* by cell ion exchange chromatography. In the example shown, two liquid cultures of cells were mixed in advance of uploading onto an ion exchange tip. Colony forming units were determined by serial dilution and plating.

**Fig. 2b.**
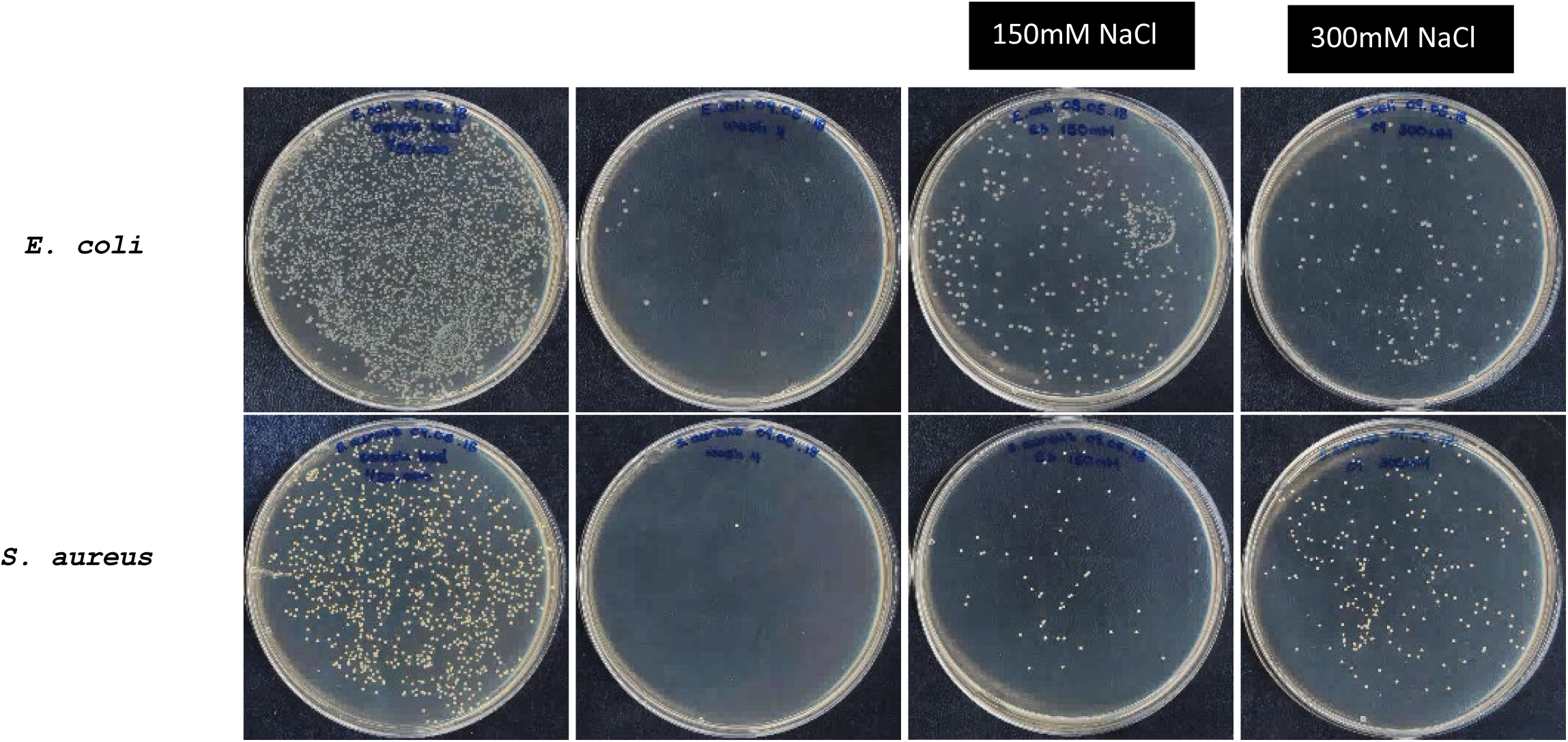
Samples from eluted fractions following independent application of *E*.*coli* and *S*.*aureus* followed by stepwise salt elution. The cross contamination of cells at 150mM NaCl is consistent with the elution profile in 2a.

These data are very similar to those obtained when separating two polypeptides differing in net surface charge at a given pH (Scopes, 1987). Further optimisation can be achieved by altering pH and by controlling the stepwise addition of elution agent, in this case NaCl, as with any ion exchange procedure. Moreover, a second dimension of separation can be achieved by using any suitable affinity resin, or a cationic resin.

In order to demonstrate that bacteria from mixed cultures can be separated by cell chromatography, two broth cultures of *E*.*coli* and *Staph. aureus* were mixed, and the combined culture uploaded onto an ion exchange column. Following chromatography and collection of the salt-eluted material, visual inspection of agar plates demonstrated that the two strains eluted at the expected salt concentrations, as observed when individual cultures were applied. For clarity, the experiment was repeated *with E*.*coli BL21* harbouring an ampicillin resistance plasmid. This makes it possible to compare the eluted fractions on LA plates with and without ampicillin. As can be seen in Figure 3, the two strains are clearly separated by a stepwise addition of sodium chloride solutions of increasing concentration.

**Figure 3.**
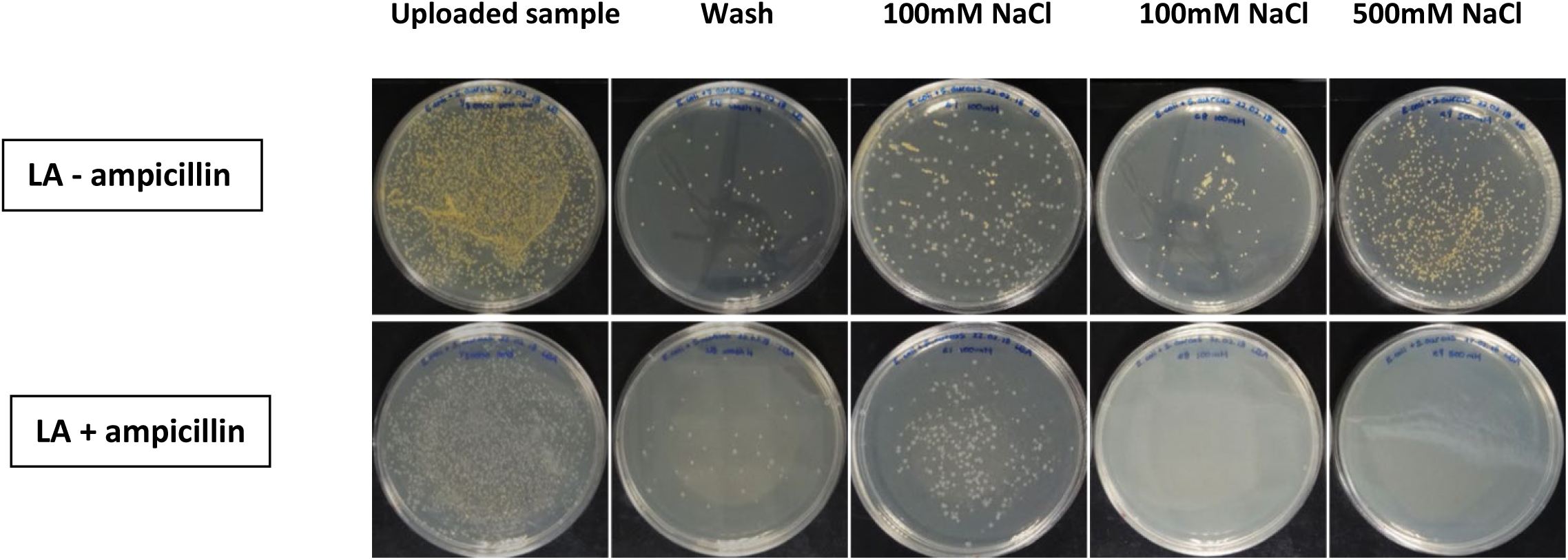
Separation of *E*.*coli* and *S. aureus* by cell chromatography. The upper series of plates show the fractions eluted from the column, followed by plating out on LA in the absence of antibiotic, where both strains grow well. In the lower gallery, the same fractions are spread onto ampicillin containing LA plates: only antibiotic resistant *E*.*coli* cells are recovered, clearly indicating that effective separation has been achieved.

### Separation of environmental strains by cell chromatography

Until relatively recently, environmental microbiologists have necessarily focused their attention on the characterisation of culturable micro-organisms. However, many microscopically observed species have remained just that until the arrival of direct genome sequencing (Venter et al, 2004). The emergence of the field of unculturable micro-organisms represents a major new opportunity for the bioprospecting. Historically, the recovery of culturable species has been the focus of taxonomic and pharmaceutical/biotechnological applications, and access to hitherto unexploited genes and metabolites represents a major shift in these fields. In Figure 4a, a series of strains isolated from a local pond were uploaded as a mixture onto a cell chromatography column. As before, a controlled application of a stepped gradient of NaCl, leads to a clear resolution of some of the species in the mixture For simplicity, strains were selected for their clear pigmentation differences, as a proof of principle for the technology. The remarkable power of resolution of cell chromatography is shown in Fig.4b, where the differential elution of the pigmented microorganisms can be clearly observed. In this experiment, an unknown set of culturable strains were separated. However, the downstream plate cultures serve to demonstrate that separation of strains has been achieved. The eluted fractions will also contain non-culturable bacteria, which can be readily accessed for many molecular and whole genome analyses.

**Figure 4.**
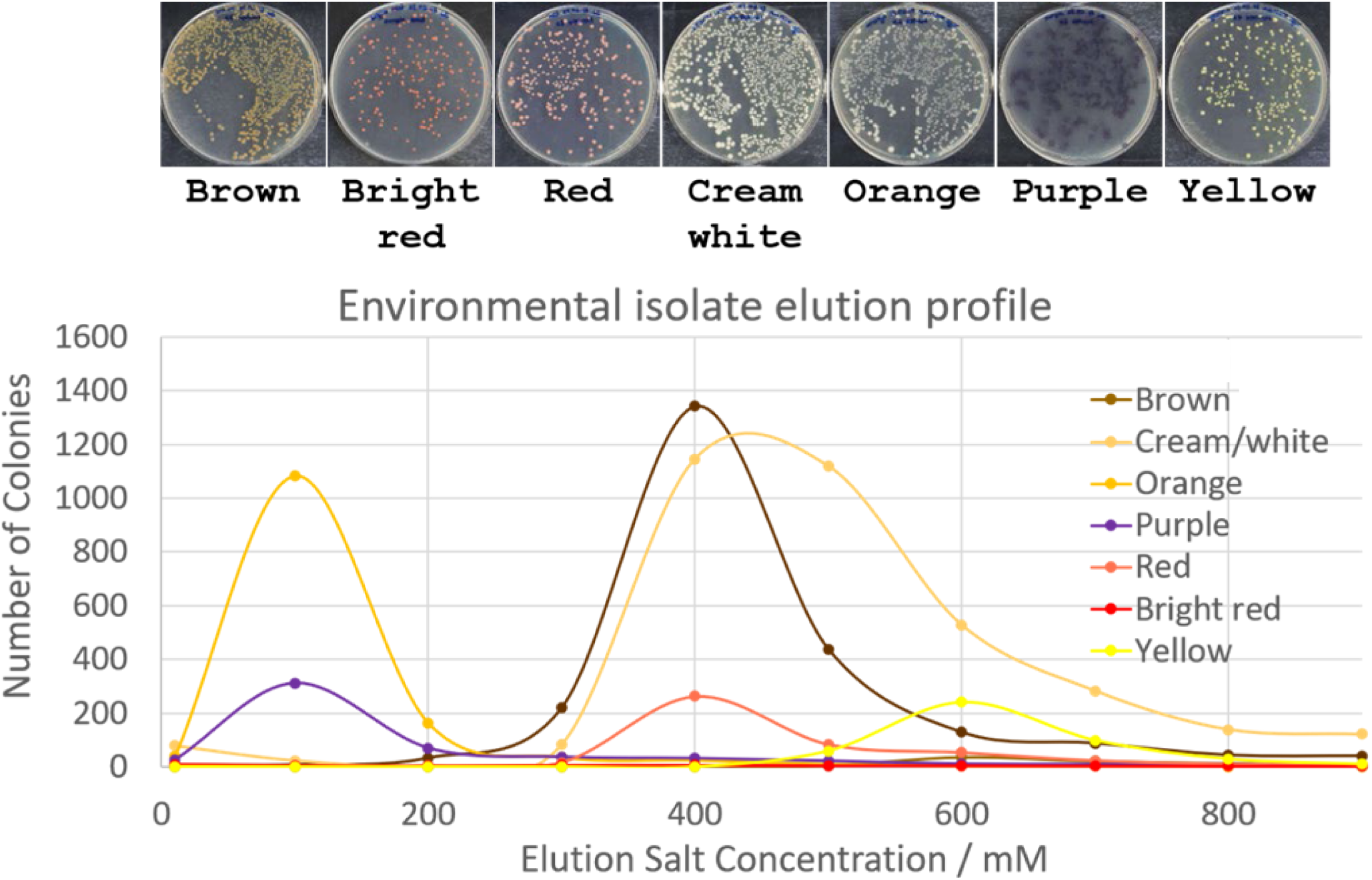
Elution profile of a mixture of pigmented strains isolated from local pond-water. A 1ml sample was uploaded directly on to a column and as in Fig. 2, eluted fractions were diluted and colony forming units counted from Petri dishes as shown in the gallery above the elution profile.

### Interrogation of immobilised bacterial cells with antibiotics

The search for novel antibiotics is a global scientific and medical priority. Ever since the serendipitous discovery of penicillin over 90 years ago, the technology used to screen antimicrobial candidates still relies on the combination of Petri dishes, solid media and cellulose disks or strips impregnated with a range of antibiotics, often at a range of concentrations. Contact with potential antibiotics indicated by a zone of growth inhibition (exemplified by Panchal et al, 2020) typically provides the first indication of antibiotic sensitivity.

*E*.*coli* BL21(DE3) cells harbouring a plasmid conferring ampicillin resistance (pET22b) were combined with an equivalent number of *S. aureus* SH1000 cells. The mixed culture was uploaded onto an ion exchange resin as described previously, followed by a number of washes with LB medium and a final short (10s) incubation of immobilised cells in LB supplemented with ampicillin at a concentration of 200μg/ml. Cells were subsequently eluted at a series of increasing concentrations of sodium chloride. Samples from all eluted fractions were subsequently plated out and incubated for 16 hours on antibiotic free Luria agar at 37°C. As can be clearly seen from Table 3, this protocol mimics the outcome of a typical plating experiment: there is an approximate 50-fold reduction in cell numbers where the strain is ampicillin sensitive. Clearly, manipulation of incubation periods, cell numbers and antibiotic concentrations would form the basis of a more thorough “interrogation”, but nonetheless, it is clear that cell chromatography protocols described here can replace traditional petri dish-based experiments in some of the fundamental procedures used in experimental microbiology. Moreover, these chromatographic protocols are much more suited to automation.

**Table 3.**
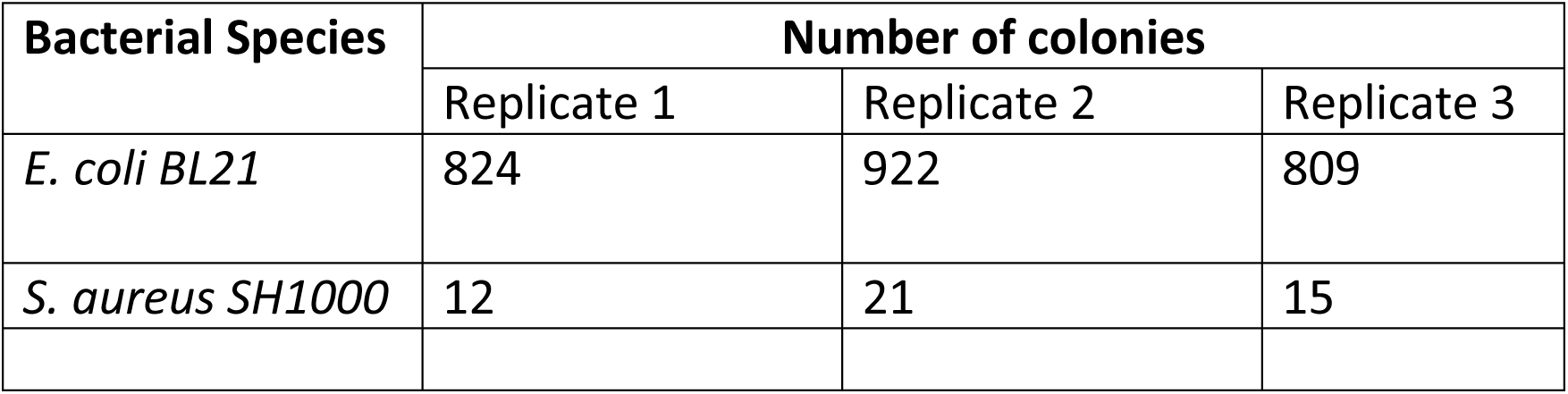
0.5ml of a suspension comprising ∼2 5000 *BL21* cells and ∼2 5000 *SH1000* cells per ml. were uploaded, followed by 25 wash steps were programmed to remove the majority of unbound bacteria. 0.5ml of LB broth containing 100µg/ml ampicillin was then applied to the column and all flow stopped for 10 seconds, after which the column was flushed out with LB. Cells were subsequently eluted from the column using 0.5ml LB broth containing 200mM NaCl and the resulting eluent plated on LB agar plates as before.

| Bacterial Species | Number of colonies |  |  |
| --- | --- | --- | --- |
|  | Replicate 1 | Replicate 2 | Replicate 3 |
| *E. coli BL21* | 824 | 922 | 809 |
| *S. aureus SH1000* | 12 | 21 | 15 |

## Discussion

Attempts to purify living bacterial cells for both analytical and preparative purposes in broth cultures have proved challenging, with most resins being utilised in a batch mode using centrifugation or magnetism to recover and interrogate resin-bound cells. While it is clearly possible to capture and separate cells that have been “fixed” in some way, such methods provide limited insight into cellular physiology. In 2006, Arvidsson et al were the first to demonstrate that cryo-gels could be used as a stationary phase for the capture and separation of viable bacteria using conventional unidirectional chromatography. Clearly, there is no *a priori* barrier to cell chromatography. Here we have utilised a novel form of capture and purification that introduces a back-and-forth mode of sample application and elution that simplifies the process and paves the way for a more controlled approach to cell analysis that goes some way to meeting the needs of high throughput, high sensitivity “omic” methods.

A population of microbial cells in an asynchronous culture (a typical batch culture) show a level of morphological diversity that is not dissimilar in principle to the diversity of polypeptides expressed in a cell. A microbial cell, or a virus particle, can be considered as a “studded” sphere (or cylinder), with the net charge distribution falling on a wide scale from negative through neutral to positive. In comparison with proteins, the considerably larger surface area of cells and viruses (or particles in general) as well as the particle size requires must be considered if retention of biological competence is required. In addition to ion exchange media, chromatography resins coupled to affinity ligands, including small molecules and antibodies, can provide a biocompatible surface for the selective purification of cells from mixed populations, thereby enriching for a specific sub-population. Such beads may be magnetic and are often incorporated into a centrifugation associated protocol. To date, however, there are very few examples of the successful chromatographic separation of cells and viruses, where the biological specimen retains full biological viability.

The column capacity of an individual column for bacteria is related to the ion exchange capacity and the equilibrium constant of the bacteria and anions in the buffer competing for the ion exchange sites. If the bacteria compete weakly, then the column capacity is low. However, if the bacteria compete strongly, then the column capacity is high, owing to the multi-valent attachment of the bacteria on the surface of the anion exchanger, where the selectivity of the bacteria is normally high, but where the kinetics of capture are relatively slow. There are several rate constants that contribute to the overall rate of capture, including those describing the binding of the bacteria to the anion exchange functional groups and the orientation of the anionic sites of the bacterial to the positive charges bound to the resin. Finally, capture of bacteria is likely to be via multi-point attachment. Since multi-point attachment increases the selectivity of binding, time is needed to maximize the attachment of a particular bacterium to the resin bead. This strong attachment is supported by back-and-forth flow through the column which gives multiple opportunities for bringing the bacterium to the ion exchange site, thereby optimising the orientation of positive and negative sites and the strongest multipoint attachment of the bacterium. However, back and forth flow is a priori deleterious to cells, damaging or killing them because of multiple chances of puncture, trapping or shearing. The columns have been designed to minimize these possible harmful interactions.

The selectivity of bacteria for ion exchangers is described by sharp isotherms as shown by the results presented in this paper. This means for any particular type of bacterial cell, of a given density and net negative charges and for any given set of buffer conditions, the bacteria are either mostly bound to the resin or mostly in solution. Furthermore, as the buffer conditions are altered there is a sharp transition of buffers where there is sorption of the bacteria to the resin compared to conditions where there is nonsorption of bacteria to the resin. Thus, small changes in the mobile phase buffer can result in a significant impact on whether a particular bacterium adheres to the column or does not. In other words, multi-point attachment makes the isotherms related to ion exchange extremely sharp, and small changes in buffer conditions (ion type and concentration) can result in large shifts in interaction affinities. In this work, a set of conditions were chosen to capture all bacteria of different types. Thereafter, the concentration of ion competing for the ion exchange sites was increased. This resulted in bacteria being released from the beads and eluted from the column. Since different bacteria have different selectivity for the anion exchanger, separation of the bacterial types has been accomplished.

The introduction of cell chromatography for the capture and interrogation of live cells in a simple chromatographic format has the potential to transform the systematic analysis of many fundamental properties of bacteria. In addition, both bioprospecting and antimicrobial discovery programs should benefit significantly from the potential for automation afforded by this technology.

